# Brain-implanted conductors amplify radiofrequency fields in rodents: advantages and risks

**DOI:** 10.1101/2022.07.20.500859

**Authors:** Mihály Vöröslakos, Omid Yaghmazadeh, Leeor Alon, Daniel K. Sodickson, György Buzsáki

## Abstract

Over the past few decades, daily exposure to radiofrequency (RF) fields has been increasing due to the rapid development of wireless and medical imaging technologies. Under extreme circumstances, exposure to very strong RF energy can lead to heating of body tissue, even resulting in tissue injury. The presence of implanted devices, moreover, can amplify RF effects on surrounding tissue. Therefore, it is important to understand the interactions of RF fields with tissue in the presence of implants, in order to establish appropriate wireless safety protocols, and also to extend the benefits of medical imaging to increasing numbers of people with implanted medical devices. This study explored the neurological effects of RF exposure in rodents implanted with neuronal recording electrodes. We exposed freely moving and anesthetized rats and mice to 950 MHz RF energy while monitoring their brain activity, temperature, and behavior. We found that RF exposure could induce fast onset firing of single neurons without heat injury. In addition, brain implants enhanced the effect of RF stimulation resulting in reversible behavioral changes. Using an optical temperature measurement system, we found greater than tenfold increase in brain temperature in the vicinity of the implant. On the one hand, our results underline the importance of careful safety assessment for brain implanted devices, but on the other hand, we also show that metal implants may be used for neurostimulation if brain temperature can be kept within safe limits.

## Introduction

The effects of radiofrequency (RF) radiation on biological tissue have been the subject of intensive research for many decades^1–3^, amplified by the advent of cellular communications. The omnipresence of Cell Towers in cities led to new urgency for understanding the effects of RF fields on the brain^4–8^. Recent and ongoing development of novel technologies expands RF applications to higher frequencies (e.g., in millimeter wave and in 5G cellular communication) and higher power levels (e.g., in high power microwave (HPM)), leading to a wider variation in public health RF exposure scenarios^9,10^. A relatively new safety concern is the effects of RF energy exposure on an increasing number of patients with implantable medical devices^11,12^. These devices generally fall into two categories. Passive devices (without powered electronic components) include joint replacements, heart valves, stents, interventional guide wires, etc. Active devices (with powered electronic components) include cardiac stimulators (pacemakers and defibrillators), neurostimulators (cochlear, deep brain, vagal nerve, and spinal cord stimulators) and the like. In recent years, we have seen a rapid increase in the number of individuals with implantable medical devices, and it is estimated that about 10% of US citizens will have a device implanted into their bodies during their lifetimes^13^.

Since RF fields are used in magnetic resonance imaging (MRI), RF effects on body tissue in the vicinity of implants have also been explored in the medical imaging literature^2,14–16^. Some implants are known contraindications for MRI, whereas others are considered MR Conditional Devices, which are defined as “item[s] that [have] been demonstrated to pose no known hazards in a specified MR environment with specified conditions of use” (American Society for Testing and Materials, F2503-20^17^; See Supplementary Table 1). The risk of RF exposure in patients implanted with metal parts can range from negligible to life-threatening^16,18^. Interactions depend on numerous factors, including the type of device, implantation geometry, length of wire leads, connectors/terminations, materials used, and more^19–26^. The assumed mechanism of interaction for most implants is between incident electromagnetic fields and conductive structures, which may produce tissue heating due to local power deposition^7,27^, may induce tissue stimulation via induced currents^26,28^ and may also result in MR image artifacts^29^. There is an ongoing interest in exploring the safety of MRI examination for patients with various kinds of new implantable devices, in order to provide these patients with the diagnostic benefits of MRI^5^. A particular case is brain implanted devices (e.g., deep brain stimulation (DBS) devices) because MRI-induced implant heating could lead to brain damage^16,18^. The risk associated with implant devices might affect an even larger number of people in the near future as there is a growing interest for commercially developed non-medical brain implants, with focus on brain-computer interfaces^13^. To prevent severe adverse events, the U.S. Food and Drug Administration requires that MRI-induced heating near implanted DBS leads be estimated using gel phantoms and computational simulations^30,31^. The focus of these guidelines is to mitigate the heating risk. However, brain tissue is excitable and MRI-induced currents in DBS devices could present additional risks for patients that can only be assessed *in vivo*. The effects of implanted metal parts exposed to RF energy are frequency-dependent, and some effects might be particular to higher frequency ranges (>10 GHz). At such frequencies, the skin depth in biological tissue is short (<1 mm). Therefore, in a person without any implants, only superficial tissues are likely to experience effects of RF energy, and deeper structures such as the brain are unlikely to be affected. The presence of a metallic implant such as a wired brain implant that extends from outside the body (or from superficial skin layers) into deeper layers, however, can transmit the effects of RF energy to deep layers, and can modulate those effects locally, e.g., by field focusing in the vicinity of sharp metallic tips.

In the work presented here, we examined the instantaneous effects of RF exposure on the neural activity in waking and anesthetized rodents. We show that the presence of a conductor in contact with brain tissue can amplify the impact of RF effects several-fold, leading to significant neurological effects.

## Material and methods

### Freely moving animal experiments

All experiments were approved by the Institutional Animal Care and Use Committee at New York University Medical Center. Animals were handled daily and accommodated to the experimenter before the surgery and recording. Mice (adult male, n = 8, C57/Bl6, 26–31 g) were kept in a vivarium on a 12-hour light/dark cycle and were housed two per cage before surgery and individually after it. Atropine (0.05 mg/kg; subcutaneous (s.c.) injection) was administered after isoflurane anesthesia induction to reduce saliva production. The body temperature was monitored and kept constant at 36–37 °C with a DC temperature controller (TCAT-LV; Physitemp, Clifton, NJ, USA). Stages of anesthesia were maintained by confirming the lack of a nociceptive reflex. The skin of the head was shaved, and the surface of the skull was cleaned by hydrogen peroxide (2%). A custom 3D-printed base plate^32^ (Form2 printer, FormLabs, Sommerville, MA) was attached to the skull using C&B Metabond dental cement (Parkell, Edgewood, NY). A stainless-steel ground screw was placed above the cerebellum. The location of the craniotomy was marked (2 mm posterior from bregma and 1.5 mm lateral to midline) and tungsten recording electrode(s) (100211, insulated with Heavy Polyimide (HML – Green), California Fine Wire Inc., Grover Beach, CA), temperature sensor (OTP-M, Opsens Solutions Inc., Canada in RF experiments and 223Fu5183-15U004, Semitec in control heating experiments) and optic fiber (200-µm diameter) were implanted into the CA1 region of the hippocampus. All devices were attached to a recoverable plastic microdrive^32^ that allowed fine positioning of the electrodes after recovery. To protect the implanted devices, a custom 3D-printed head cap was attached to the base plate^33^. Each animal recovered for at least 7 days prior to electrical recordings. Ketoprofen was administered up to 2 days after surgery (0.5 mg/mL, s.c. injection).

The collected data was digitized at 20 kSample/s using an RHD2000 recording system (Intan technologies, Los Angeles, CA). A baseline session and radiofrequency stimulation session(s) were recorded from each mouse. The duration of the sessions depended on whether there was an afterdischarge/behavioral response or not. The behavior of the animal was monitored with a camera (Basler ACE2, #a2A2590-60ucBAS, at 25 frames per second), that was placed above the home cage. A mirror was placed in front of the home cage providing a side view of the animal behavior.

### Urethane-anesthetized experiments in rats

Adult, male rats (n = 2, Long Evans, 350-450 g) were used for acute experiments. Rats were implanted with silicon probe (Buzsaki 5×12, NeuroNexus, Ann Arbor, MI) under urethane anesthesia (1.3–1.5 g/kg, intraperitoneal injection). A custom designed, 3D-printed plastic base was attached to the skull and a silicon probe was implanted into the somatosensory cortex (18-degree angle) using a microdrive system^32^. The rat was placed inside a Transverse ElectroMagnetic (TEM) cell (up to 2 GHz, 50 ohm characteristic impedance, and 36 cm x 48 cm x 15 cm cavity size). We waited 60 minutes before data collection. Baseline recording (40-60 min) was performed before RF stimulation which was applied using 3 different intensities (2 s stimulation followed by a 2 s stimulation-free epoch, at least 250 trials, 950 MHz, 6.5, 10, and 41 W power). After RF stimulation, post-baseline recording was performed (40-60 min). The collected data was digitized at 20 kSample/s using an RHD2000 recording system (Intan technologies, Los Angeles, CA).

### Urethane-anesthetized experiments in mice

Adult male mice (n = 5, C57Bl6, 26-33g) were anesthetized with urethane (1.3–1.5 g/kg, intraperitoneal injection). 3 out of 5 mice were implanted in the hippocampus with a triplet of tungsten wires (50 µm diameter, 7.5 cm long, insulated with Heavy Polyimide, 100211, California Fine Wire Inc., Grover Beach, CA) accompanied with an optical temperature probe (0.3 mm in diameter, PRB-100-01M-STM Osensa Innovation Corp.) which does not absorb RF and was attached to a custom-made microdrive^32^. The temperature sensor was fixed to the body of the microdrive while the wire triplet was attached to its movable shuttle. Temperature recordings were collected from the corresponding signal conditioner (FTX-300-LUX+, Osensa Innovative Corp.). Animals were exposed to RF from a nearby patch antenna. When the wire triplet was inside the brain, 3 or 4 short pulses (0.5 s) of RF energy with varying input power to the antenna (9-68 W, measured by a power meter (U2001A Power Sensor connected to N9912A Field Fox Spectrum analyzer, Agilent/Keysight) and a bidirectional coupler (778D, Agilent/Keysight)) were applied. Afterward, the wire triplet was withdrawn from the brain (with the temperature probe staying in place) and the same procedure was repeated. 50 s RF pulses were applied to induce measurable heating in the absence of metal electrodes in the brain (n = 4 mice). In n=2 of these mice we also changed the location of the portion of the wire triplet that was outside the animals’ brain relative to the antenna and recorded temperature responses to short (0.5 s) RF energy pulses of the highest input power (68 W). In an additional n=2 mice only the temperature probe was implanted and temperature responses to 50 s long pulses were recorded. In a single male mouse, only a wire triplet was implanted in the hippocampus. The outside part of the wire was placed close to the patch antenna to induce the highest temperature rise in the brain. A set of 4 short (0.5 s) RF energy pulses of the highest input power (68 W) were applied. Then the animal was euthanized, and the brain was collected for histology to observe local damage to the brain tissue.

### Calculation of RF-induced temperature rise

As the brain temperature *in-vivo* is constantly fluctuating^34,35^, the baseline for temperature measurement when an RF pulse is applied is often not flat. To account for the pre-stimulus temperature changes, the effective RF-induced temperature rise (ΔT) was calculated by the magnitude of temperature change during the RF-On period (ΔT_1_) minus the temperature change during a similar period of time in the baseline right before the stimulation onset (ΔT_2_; ΔT_=_ΔT_1-_ ΔT_2_)^36^. If the temperature changes during the pre-stimulus period were not linear that stimulation pulse was discarded (about 5 % of all pulses).

### Radiofrequency stimulation

Two different antennas were used in our experiments: a TEM cell (TBTC3, Tekbox Digital Solutions, Vietnam) to create uniform electric fields in our initial experiments, and a patch antenna (M3070100P11206-B, Vente/TerraWave) that allowed more flexible experimental design (e.g., using a camera to monitor animals’ behavior, changing the orientation of the antenna relative to the animal). The TEM cell was used in the urethane-anesthetized rats, and the patch antenna in awake mice and in urethane-anesthetized mice experiments. Each antenna was driven by an ultra-low noise, dual channel RF signal generator (SynthHD, Windfreak Tech) with a maximum power level of 0.1 W. To achieve higher power levels, we inserted an amplifier (ZHL-100W-13+; Mini-Circuits) between the signal generator and the patch /TEM cell^36^.

### Single unit analysis

A concatenated signal file was prepared by merging all recordings from a single animal from a single day. To improve the efficacy of spike sorting, stimulation induced onset and offset artefacts were removed before automatic spike sorting (10 ms before and 100 ms after the detected artefacts, linear interpolation between timestamps). Putative single units were first sorted using Kilosort^37^ and then manually curated using Phy (https://phy-contrib.readthedocs.io/). After extracting timestamps of each putative single unit activity, peristimulus time histograms and firing rate gains were analyzed using a custom MATLAB (Mathworks, Natick, MA) script.

### Brain state scoring

Spectrograms for brain state scoring were constructed with a 1 s sliding 10 s window fast Fourier transform of 1250 Hz local field potential (LFP) data at log-spaced frequencies between 1 and 100 Hz. Three types of signals were used to score state in our recordings: broadband LFP, narrowband theta frequency LFP, and electromyogram (EMG). For broadband LFP signal, principal components analysis (PCA) was applied to the z-transformed (1-100 Hz) spectrogram. The first principal component in all cases was based on power in the low (< 20 Hz) frequency range and had oppositely weighted power in the (> 32 Hz) higher frequencies. Theta dominance was taken to be the ratio between the power of 5-10 Hz and 2-16 Hz bands from the spectrogram. EMG was extracted from the intracranially recorded signals by detecting the zero time-lag correlation coefficients I between 300-600 Hz filtered signals (using a Butterworth filter at 300 – 600 Hz with filter shoulders spanning to 275 – 625 Hz) recorded at all sites^38^. The state scoring algorithm was performed by a series of divisions with thresholds set at the trough between the peaks of distributions in these three metrics^39^ (https://github.com/buzsakilab/buzcode/tree/master/detectors/detectStates/SleepScoreMaster).

After automated brain state scoring, all states were manually reviewed by the experimenter and minor corrections were made when discrepancies between automated scoring and user assessment occurred^39^.

### Behavioral observations during RF stimulation

The video was scored during and after RF stimulation. During induced seizures, scoring was based on the Racine scale^40^, with the following stages: (0) no change in behavior; (1) facial movements; (2) head nodding; (3) forelimb clonus; (4) rearing; (5) rearing and falling (generalized motor convulsions). In experiments when no seizure was expected, for example, with screw electrodes placed above the cerebellum, overt change in behavior, such as sudden motion, locomotion, head turns and sniffing or head scratching within 15 s of RF exposure were regarded as potentially “evoked”.

### Statistical Analysis

Statistical analyses were performed with MATLAB built-in functions or custom-made scripts. The unit of analysis was typically identified as single neurons. In a few cases, the unit of analysis was sessions or animals, and this is stated in the text. Unless otherwise noted, non-parametric two-tailed Wilcoxon rank-sum (equivalent to Mann-Whitney U-test) or Wilcoxon signed-rank test was used. On box plots, the central mark indicates the median, bottom and top edges of the box indicate the 25^th^ and 75^th^ percentiles, respectively, and whiskers extend to the most extreme data points not considered outliers. Due to experimental design constraints, the experimenter was not blind to the manipulation performed during the experiment (i.e., RF-stimulation).

## Results

### RF-induced fast onset firing of neurons

To address the possibility that RF exposure can change neural activity, we implanted multi-shank, multi-site silicon probes^32^ into the somatosensory cortex of anesthetized rats (n = 2). These experiments were performed inside a TEM cell antenna (Fig. 1a,b). Sessions consisted of baseline recording (30 min stimulation free period) and intermittent RF radiation (2 s RF stimulation followed by 2 s stimulation free periods, at least 250 trials; Fig. 1a,b). Although transient RF-induced artifacts (at the stimulation onsets and offsets) were present in the LFP, they did not affect the waveform or amplitude of the extracellularly recorded action potentials (‘spikes’)^36^. As Fig. 1c,d illustrates, RF stimulation (6.5 W to 41 W) induced either excitation (n = 23 single neurons) or suppression (n = 30 single neurons) in a subset of the recorded population (n = 53 single neurons). The magnitude of both excitatory (ΔFR: 1.27% ± 0.18, 1.61% ± 0.3 and 3.84% ± 0.65 for 6.5, 10, and 41 W, respectively; n = 23 single units out of 53) and inhibitory (ΔFR: 0.15% ± 0.36, -1.31% ± 0.4, -5.5% ± 0.98 for 6.5, 10, and 41 W, respectively; n = 30 single neurons out of 53) effects was proportional with the strength of the RF exposure (Fig. 1e). The onset latency was rapid (around 100 ms), and the effect (either suppression or excitation) was maintained throughout a single simulation epoch and across trials (n = 250 trials).

**Figure 1.**
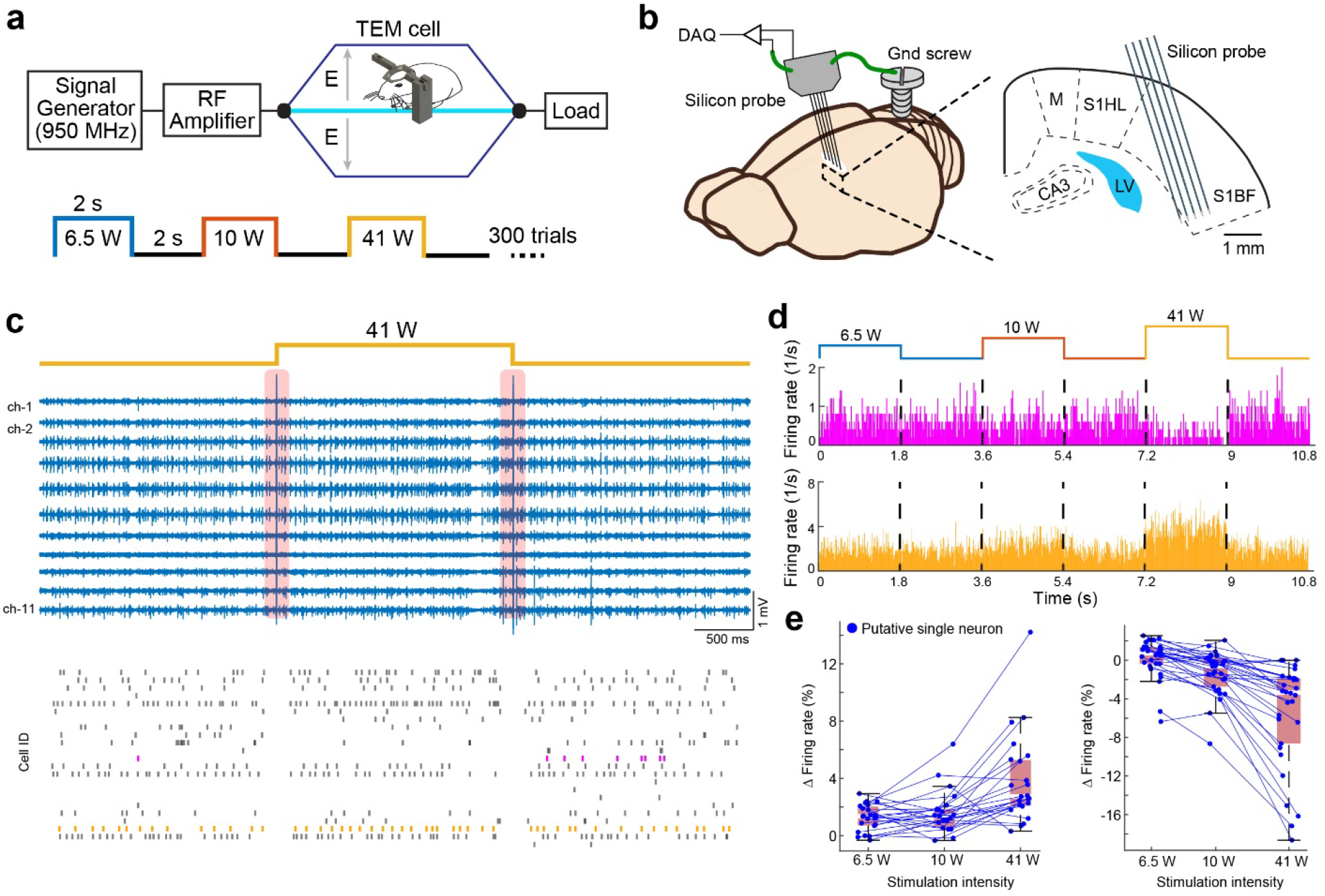
RF stimulation induced modulation of single unit firing in anesthetized rat. **a** Schematic of experimental setup. Urethane-anesthetized rat was placed inside a TEM cell antenna and 950 MHz RF-stimulation was applied. **b** Left: schematic of recording configuration. Silicon probe was used to record the neural activity. Stainless-steel ground screw was placed above the cerebellum and was connected to the recording preamplifier. Right: brain schematic shows the location of the recording electrode in the somatosensory cortex. **c** Electrophysiology signal was not affected by RF stimulation (high pass filtered traces are shown from one shank during 41 W RF stimulation), except for the induced turn on/off artifacts (red rectangles indicate the removed data, more details in Methods). Raster plots show the response of 14 putative single units recorded from channel 1-4. Note, the reduced spiking of the cell in magenta and the increased spiking of the cell in orange. **d** Response of example neurons for RF stimulation (6.5, 10 and 41W input power to TEM cell was used, 2 s stimulation was followed by 2 s control periods for at least 250 trials). Activity of one neuron was suppressed (in magenta, top) while the other elevated (in orange, bottom) by RF stimulation in an intensity-dependent manner, as shown by their peristimulus time histograms. **e** RF stimulation significantly modulated the spiking activity of cells (n = 23 cells increased and n = 30 cells decreased their spiking activity, p < 0.05, Wilcoxon rank-sum test). Box plots are shown in the background for each stimulation intensity (red rectangles).

### RF stimulation of neuronal tissue in presence of conductor

In this experiment, we observed that high-power RF stimulation (60 W, 950 MHz), when applied to a patch antenna placed underneath the homecage of chronically implanted mice with tungsten electrodes or silicon probes with 3 or 64 recording sites, respectively, occasionally induced epileptic afterdischarges in the hippocampus (Fig. 2a-c). These discharges were very similar to those induced by repetitive electrical stimulation of the hippocampus or surrounding structures^40^. Changing the frequency of the stimulation (100 Hz instead of 8 Hz) did not affect the occurrence of afterdischarges (Suppl. Fig. 2). The afterdischarge was followed by postictal depression and secondary afterdischarges (Fig. 2c bottom), as well as head nodding and occasional ‘wet-dog shakes’, typical of kindled seizures^40,41^ (n = 13 sessions in 6 mice, Suppl. Video 1). In contrast, the same RF exposure had no detectable behavioral effects in non-implanted mice (n = 10 sessions in 7 mice; Table 1, Suppl. Video 2). Our electrical recording setup had multiple conductor elements, including wire or silicon recording probes in the brain, stainless steel screw in the skull that served as ground/reference electrode, wires connecting to the head-stage preamplifiers, the preamplifier itself, the copper protector shielding and a coaxial cable routing the brain signals to farther amplifiers (Fig. 2a). We reasoned that any of these components or their combinations can contribute to local RF-amplification and, therefore, we systematically examined the configurations which did or did not induce behavioral effects. In several cases, the same mouse was repeatedly tested while we reconfigured the head-stage by disconnecting or removing components, while the electrodes were fixed inside the brain (Fig. 3b-h; Table 1).

**Table 1.** Number of sessions and animals used to investigate the role of metallic components. (related to Figure 3).

| Condition | # of sessions/animals | Response | Notes |
| --- | --- | --- | --- |
| a | 10/7 | No | None |
| b | 8/4 | No | None |
| c-d | 11/4 | No | None |
| e-f | 4/4 | Yes, in 1 session | Head scratching |
| g-h | 9/5 | Yes, in 7 sessions | Circling behavior, head scratching, running and jumping |

**Figure 2.**
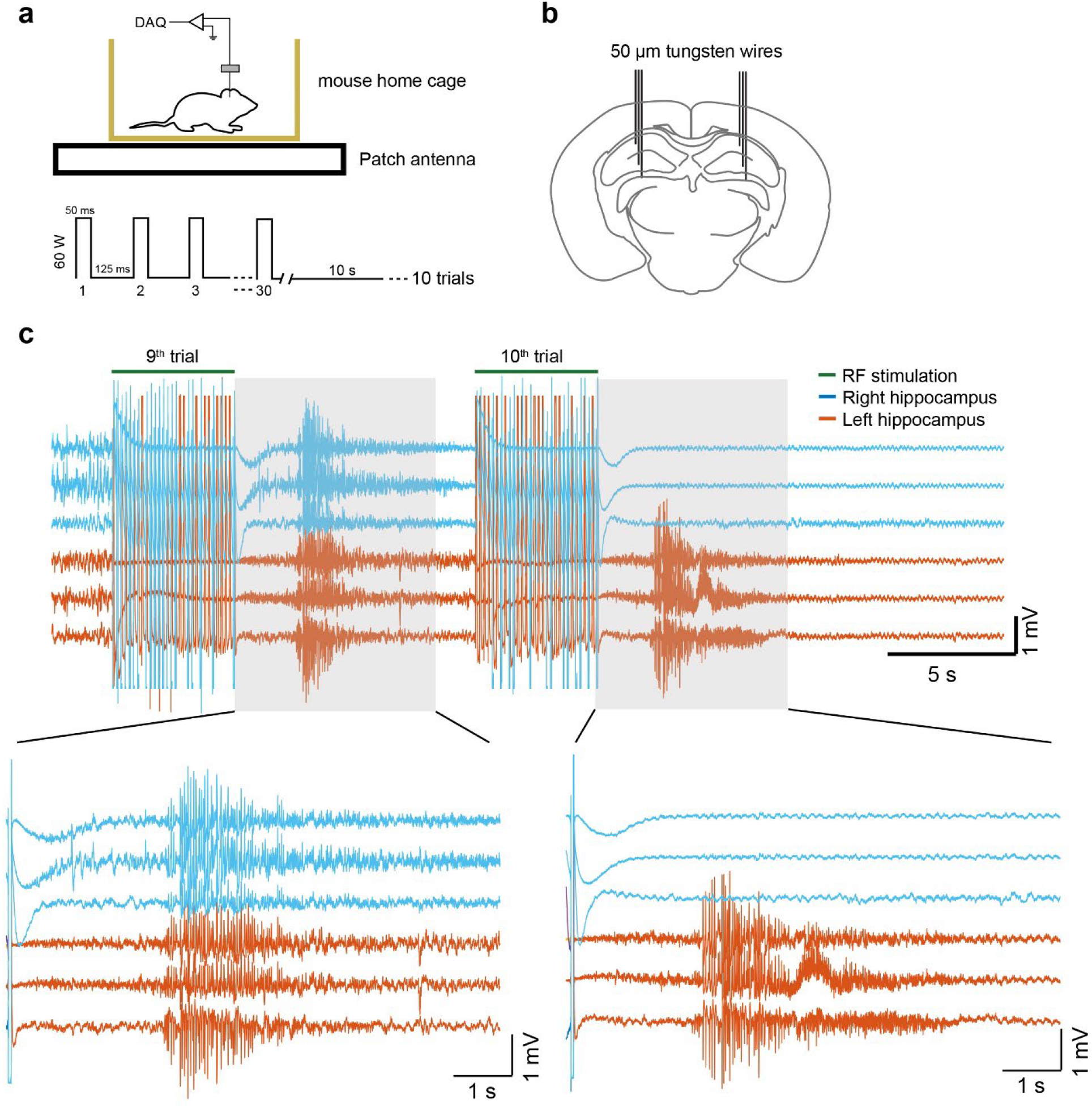
RF-induced epileptic seizures. **a** Schematic of the experimental setup. An implanted mouse was placed in its home-cage on top of a patch antenna (top) and 950 MHz RF stimulation was applied (50 ms pulses at 8 Hz, 30 repetitions in a trial, 10 trials at 60 W input power; bottom part). **b** Tungsten wire triplets (50 µm wires) were implanted in both hippocampi in a wild type adult mouse. **c** RF stimulation induced epileptic patterns after repeated stimulation. Bilateral or unilateral effects are observed after the 9^th^ and 10^th^ trains, respectively.

**Figure 3.**
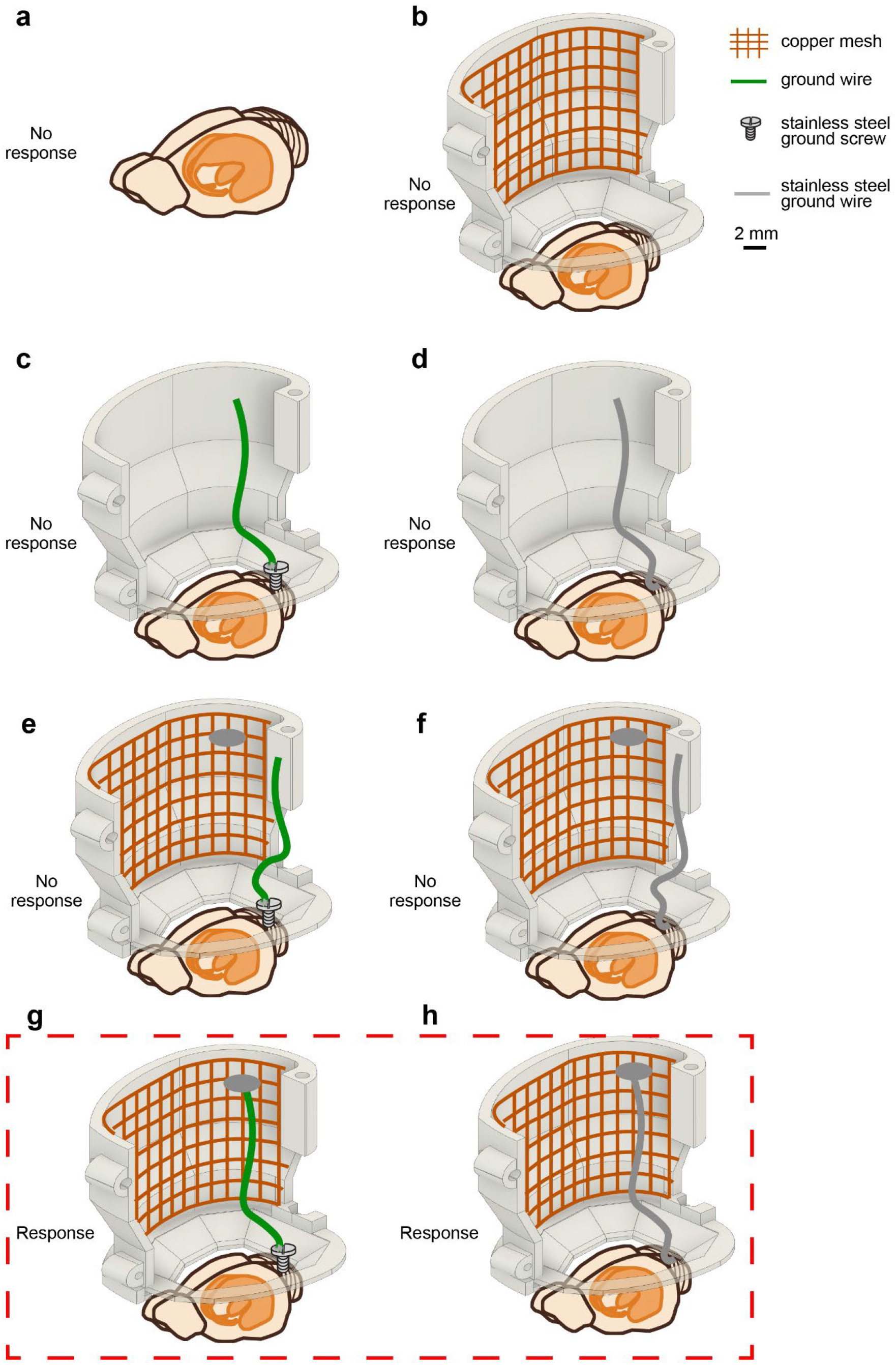
Role of metallic components in RF induced neuronal effects. Mice were tested in their home cage with a patch antenna placed under the cage (as in Figure 2a). In each condition RF stimulation was applied. **a** Non-implanted animal. **b** Head-cap with copper mesh. **c and d** Experiments with ground wires only. **c** Stainless steel screw with a copper wire (green, 100 µm in diameter). **d** Stainless-steel wire inserted into the cerebellum (100 µm in diameter). **e and f** Same configuration as **c** and **d**, with copper mesh added to the inside of the plastic cap. **g and h** Same as **e** and **f** but the ground wire soldered to the copper mesh shielding. Only the bracketed configurations (**g** and **h)** resulted in behavioral effects in all animals (see Table 1).

Removing the cable and the head-stage did not prevent induction of the observed behavioral effects of RF stimulation, excluding the sole contribution of some ground loop or artificial current injection into the brain by the preamplifier due to RF interference with electronic components. In animals which had no direct metal connection with the brain or skull, no behavioral effects were detected even though they had a large copper mesh shield protector on their head (Fig. 3c). This result reduced the possibility that steering of the RF beam was achieved due to field enhancement by the sole presence of the wire mesh. On the other hand, when the copper mesh protector was connected to brain tissue via a stainless-steel wire or through a piece of copper wire connected to a stainless-steel screw in the skull, behavioral effects, including circling, head scratching, running, and jumping, were consistently induced (Fig. 3e, f; Suppl. Video 3). The presence of the short ground wire and skull screw was not sufficient to induce behavioral effects reliably and consistently (Fig. 3d, e; Table 1). Furthermore, when both the ground wire and copper mesh shield was present but unconnected, in three of the four sessions no behavioral consequences were induced by RF stimulations. In the remaining case, RF stimulation elicited head scratching. These observations suggest that the metal connection with the brain is a critical component of the RF-induced neuronal effects (Fig. 3g, h; Table 1).

### Brain heating induced by radiofrequency stimulation

A well-documented biological effect of RF (microwave) stimulation is its heat production in biological tissue^42^. To monitor brain temperature changes, we used a metal-free optical temperature measurement system to avoid interference of RF energy with electronic components^36,43^. The tip of the fiber-optic temperature probe (1.1 mm diameter) was implanted in the brain. The impact of RF stimulation on brain temperature changes was tested in freely moving mice with implanted wires in the hippocampus (Fig. 4a,b). Like the experiments illustrated in Fig. 2, RF stimulation consisted of 50 ms pulses followed by 125 ms no stimulation, 60 repetitions, 60 W input power, followed by 10 s no stimulation (recovery) epochs (Fig. 4c). The stimulation protocol induced both short-lasting and cumulative effects. During each RF stimulation train, brain temperature increased rapidly by approximately 1.5 °C within 10 s from stimulation onset, followed by a rapid but not complete recovery (Fig. 4d). As a result, repetitive trains brought about a cumulative ∼ 6 °C increase of brain temperature. When the orientation of the patch antenna relative to the head was changed by 90°, the impact of RF stimulation was dramatically changed (Fig. 4d). Occasionally, RF stimulation in animals with implanted electrodes elicited an afterdischarge (Fig. 5a), as in the drug-free mice. The evoked afterdischarge did not have a clear temperature threshold (Fig. 5b). In addition, optically induced heat was not able to generate afterdischarges or any behavioral effects in a freely moving mouse (Suppl. Fig. 3), suggesting that the underlying mechanism to evoke afterdischarges is not solely driven by the absolute value of brain temperature.

**Figure 4.**
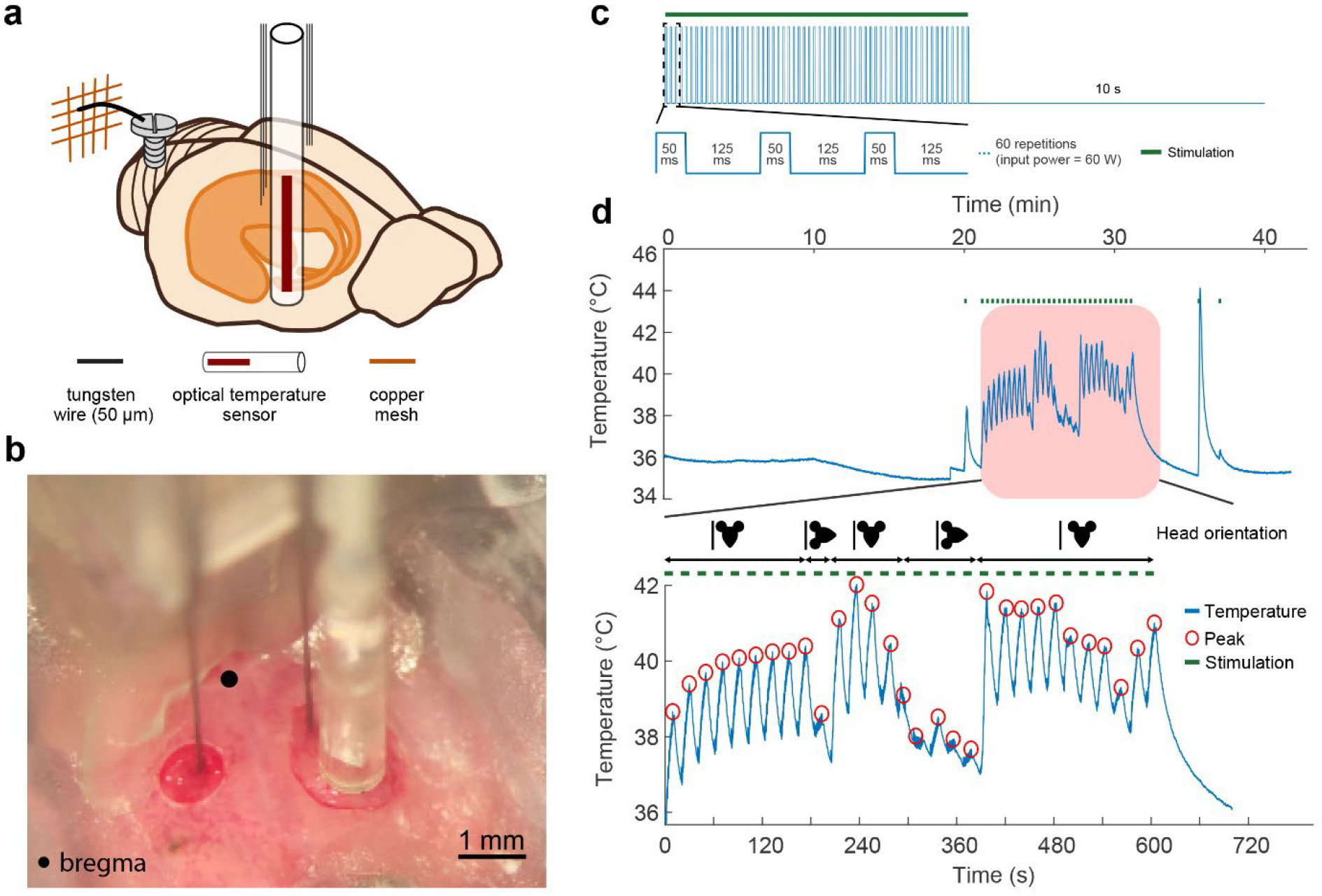
RF-induced heating in the wire implanted mice brain. **a** Mice were implanted with tungsten triplet electrodes bilaterally in the CA1 region of the hippocampus and unilaterally with the optical temperature sensor. Stainless-steel screw was used as ground and was attached to the protective copper mesh. **b** Intraoperative photograph shows the recording electrodes and temperature sensor. **c** Radiofrequency stimulation schedule. Mice in their home cage were placed on top of a patch antenna (as in Fig. 2a). RF stimulation was applied with the following parameters: 50 ms pulses at 8 Hz, 60 repetitions, 60 W input power. **d** Top: brain temperature recorded with the optical temperature sensor before, during and after RF stimulation. Note the RF-induced repeating temperature rises. Bottom: zoomed portion of the trace. The head orientation of the mouse was changed (schematic above the trace shows the orientation relative to the patch antenna), which changed the RF-induced heating.

**Figure 5.**
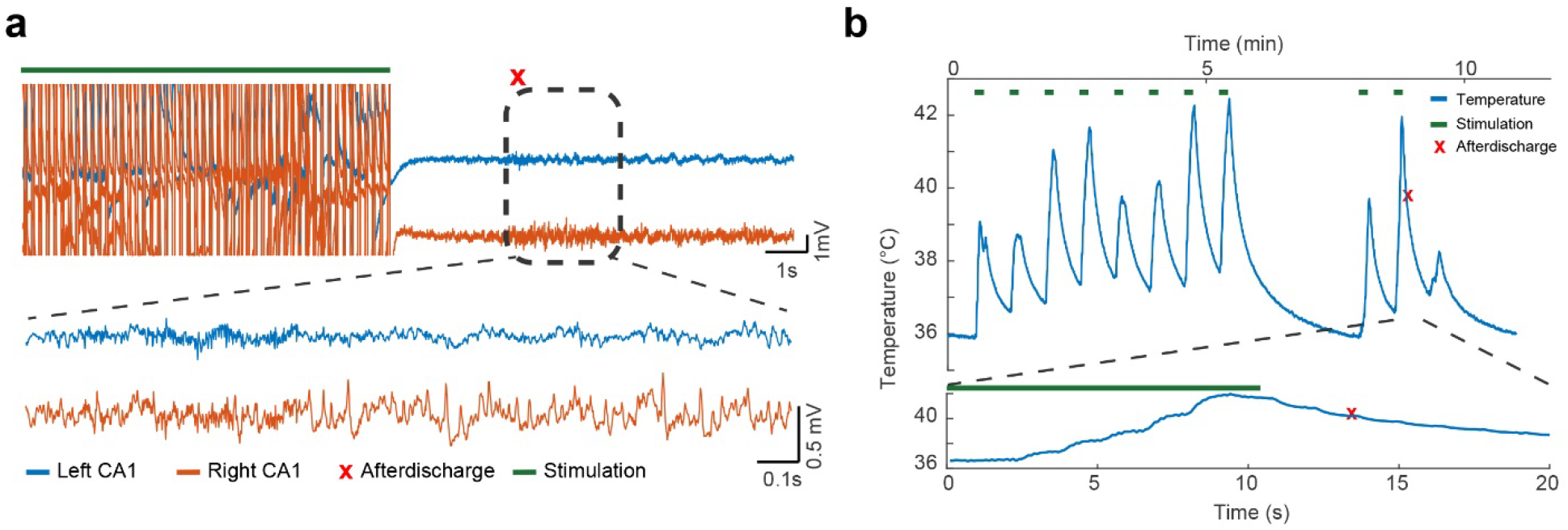
Slow RF-induced temperature rise alone cannot explain afterdischarge generation. **a** Brain temperature over time during RF stimulations (patch antenna, 50 ms pulses at 8 Hz, 60 repetitions, 60 W input power). Bottom trace shows a zoomed-in trial during which an epileptic afterdischarge was observed. **b** Electrophysiological activity during and after RF stimulation recorded bilaterally in the hippocampus (left and right hippocampal CA1 recording sites). Note that the seizure was mainly localized to the right hippocampus.

### Metal in the brain enhances RF-induced heating

We directly measured and compared induced temperature changes in the presence and absence of metal wires in order to quantify the role of conductive electrodes in the brain. To monitor brain temperature changes, we used a smaller diameter (0.3 mm) optical temperature probe. A 3D printed head-cap was mounted on the animal’s skull to allow head fixation of the mouse under urethane anesthesia and RF stimulation applied by a nearby patch antenna (Fig. 6a). A triplet of tungsten wire electrodes (50 µm in diameter) together with the optical temperature probe, both attached to a microdrive (500-800 µm distance between the wires and the probe; Fig. 6a), were implanted in the hippocampus. The temperature probe was fixed to the microdrive’s body while the wire triplet was attached to its moveable shuttle. Applying short (0.5 s) RF pulses of different power intensity (9 to 68W input power to the antenna), we observed few degrees to tens of degrees of transient increase in brain temperature. After this first stage of the experiment, the metal electrodes were withdrawn from the brain, and we applied the same RF energy continuously for a longer period (50 s) to ensure a clear reading of temperature rise and measured the brain temperature in the absence of metallic wires. Despite the longer RF energy deposition, brain temperature increased by ≤1.3 °C in the absence of wires, demonstrating the large impact of the metal electrodes on both the magnitude and the speed of temperature changes. Fig. 6b shows the temperature rise during the applied RF pulse, with highest intensity in our experimental preparation (68 W input power into the antenna), in the cases of presence and absence of the metallic wires in the brain, respectively. Note that a much shorter (0.5 s vs 50 s) RF pulse of the same intensity induces a 30-40-fold higher temperature rise when wires are present in the brain. The observed heat effect was dose-dependent both in the presence and in the absence of metal wires (Fig. 6c). Because the pulse duration for the two cases was different, a more direct comparison would require the evaluation of the rate of the temperature rise (dT/dt), due to RF exposure, rather than the induced temperature rise (ΔT). Fig. 6d shows that the rate of temperature increase is more than three orders of magnitude higher in the presence of the wires in the brain. Next, we examined whether the position of the non-implanted portion of the wire triplet relative to the antenna (hence the quality of its coupling with the radiated RF fields and the amount of energy it absorbs) has an impact on the temperature increase in the brain. Fig. 6e shows that the position of the wire relative to the antenna has an important effect on the temperature rise in the brain with wires closer to the antenna inducing higher temperature rise^44,45^.

**Figure 6.**
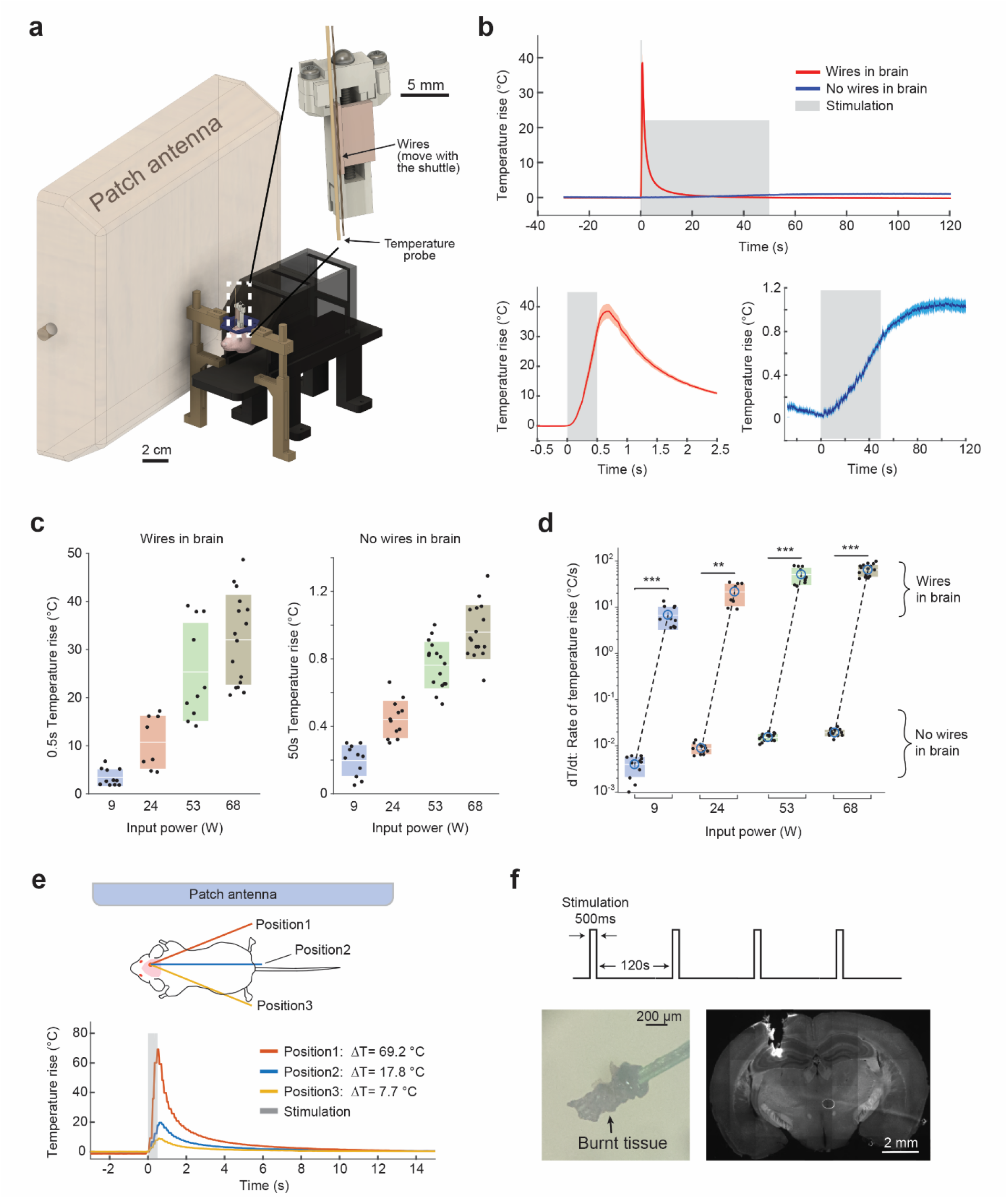
RF-induced temperature rises in presence and in absence of implanted metallic electrodes in the brain. **a** Schematic of experimental setup. Implanted mouse attached to a head-fixation set-up is placed next to a patch antenna in a fixed relative position. A triplet of 50 µm tungsten wires and a 300 µm wide optical temperature sensor are attached to a microdrive (white dashed box). Right: the temperature sensor is fixed to the body of the microdrive, but the wires can be moved with its shuttle. **b** RF-induced temperature rises in mouse brain in-vivo exposed to highest RF input power to the antenna in our set-up (68 W) in presence of the wire triplet in the brain (n=3 mice) and with no implanted wires (n=4 mice). Bottom: zoomed-in temperature responses. Note the different time course between the two conditions. Grey shading shows periods when RF radiation is applied. **c** RF-induced temperature rises in presence (left) and absence (right) of a wire triplet for different RF power levels. Note that the exposure time is different for the two cases. **d** Comparison of the rate of the temperature rise during RF stimulation periods for presence versus absence of implanted wires in the brain (same data as in **c**). Dashed lines connect the mean values of the rate of the temperature rise in presence versus absence of implanted wires for each RF input power intensity. Note the logarithmic scale on the y-axis. **e** The magnitude of the induced temperature rise depends on the position of the external portion of the wire triplet relative to the antenna. Schematic illustration of the experimental preparation (top). The closer the wire is placed to the surface of the antenna the higher the temperature rise in the brain (RF input power of 68W is applied). Gray shading shows the RF stimulation period. **f** Example of permanent damage of the brain tissue surrounding the wire triplet after four 500-ms long (120 s apart) pulses of 68 W input power RF energy (with wires in position-1 as in **e**. Schematic of RF stimulation paradigm. Burnt tissue attached to the extracted wires from the brain (bottom left). Histological image of the brain at the implant location (bottom right).

Finally, we examined whether strong RF pulses could bring about a non-reversible damage to the brain tissue surrounding the implanted wires (Fig. 6f). Indeed, a mere set of four 0.5-second-long RF pulses at the highest intensity (68 W) with the wire triplet positioned close to the antenna (position1 in Fig. 6e) were sufficient to induce tissue burning around the implanted wire triplet (Fig. 6f).

## Discussion

We examined how metal conductor in contact with brain tissue enhanced the effect of RF stimulation and found that even a standard recording electrode can induce more than a tenfold increase in local temperature. Moderate levels of RF energy exposure reliably affected firing rates of neurons, possibly through local heat-enhancement by the electrodes.

### Electrophysiological and behavioral observations

Electrophysiological experiments we performed prior to the work reported here, using thin wires or metal-containing silicon electrodes, showed that RF stimulation can reliably induce action potentials or suppress spontaneously occurring action potentials in a fraction of cortical neurons, comparable to the fractions reported in transcranial electrical stimulation experiments^46,47^. The induced effects were observed 100 ms after stimulation onset and were sustained throughout the stimulation. The magnitude of the effect was proportional to the deposited RF energy intensity. Using relatively moderate RF power levels, RF-induced changes in neuronal spiking activity were maintained across hundreds of stimulation trials, indicating that neurons were not damaged by weak RF stimulation. Repetitive stimulus trains of higher intensity (60 W) in the theta frequency range (6 Hz), could induce hippocampal epileptic afterdischarges, accompanied by behavioral manifestations, including characteristic postictal head and body shaking (“wet-dog” shaking), rearing, and falling. These findings confirm and extend previous electrophysiological experiments in vitro^48–50^. They also support previous EEG studies in both humans and experimental animals that exposure to weak electromagnetic fields, comparable to those emitted by cell phones, can influence brain electrical activity^51–55^. However, when non-implanted animals were exposed to identical RF power and frequency, we did not observe any reliable behavioral effects. Therefore, we tentatively concluded that in our experiments and possibly all previous experiments using metal electrodes, the presence of the conductive metal amplified the RF effects. This conclusion was supported by our subsequent set of experiments, in which we systematically varied the configuration of the recording head stage. Metal above the head and not in direct contact with the skull or brain surface did not affect behavior. In contrast, when either a wire or a metal screw was in contact with brain tissue, behavioral manifestations and afterdischarges could be detected.

### Metal electrodes enhance RF-induced effects

Using optical temperature measurements allowed us to directly measure interference-free RF-induced heat effects. In the intact brain, moderate-intensity RF stimulation induced only a few degrees change of temperature which remained within the range of physiological temperature variation^35,56^, although the RF exposure was above the levels recommended by regulatory agencies. (US regulations limits, set by the Federal Communication Commission^57^ are specific absorption rate [SAR] values of 1.6 W/Kg and 8 W/Kg, averaged over 1 gram of tissue, for public and occupational/controlled cell phone exposures, respectively.) SAR value was about 29 W/Kg in our head-fixed set-up applying 37.5 W input power into the patch antenna^36^.

Our measurements also revealed that the mere presence of a short wire in the brain and above the skull could enhance local temperature by several tens of degrees Celsius within just few hundreds of milliseconds. It should be noted that our temperature measurement was obtained several hundred µm from the metal wire, thus it is likely that the temperature increase in the immediate vicinity of the wire was even higher and steeper. Such large and sudden changes of local heat was sufficient to affect neuronal activity, provided that the product of the magnitude and duration of RF stimulation remains below a certain threshold. Yet, without metal-amplification RF radiation (with in-situ electric fields up to 230 V/m) does not impact neuronal activity, as demonstrated by a recent in vivo Ca^2+^ imaging experiment^36^. Overall, our experiments indicate that RF effects on excitable tissue are mediated primarily by transient or sustained heat. Yet, we cannot conclusively exclude previously postulated non-thermal effects of RF irradiation, such as vibration-mediated thermo-elastic effects in brain tissue^58–60^.

### RF-induced risks associated with implantable devices

Patients with brain implants are potentially vulnerable to moderate and high levels of ambient RF radiation. The strong metal amplification effect we observed is consistent with concerns about risks to patients with metallic implants during MRI examinations^30,61–63^ or other occasions of exposure to high intensity RF energy. Our experiments emphasize that the risks are not only limited to tissue damage at higher levels of RF exposure, but they extend to undesired modulation of the ongoing neural activity in the vicinity of the implanted metal parts even at moderate or lower levels of RF exposure. This is particularly important as there is a growing interest in non-medical brain implants^64^. While safety guidelines related to RF exposure are defined for non-implanted subjects^57^, and while implants are generally subjected to careful scrutiny for safety in MRI settings, patients with metallic brain implants could be at risk in some real-life scenarios of RF exposure. In particular, individuals with implanted metal devices may be vulnerable to novel military or civil high-power microwave applications (e.g., base stations or high-powered Wi-Fi routers).

### RF-powered brain stimulation implants

While our findings call attention to potentially damaging consequences of RF exposure, they also suggest an opportunity to explore RF stimulation as a non-invasive method to affect brain activity. Weak RF stimulation excited or inhibited a considerable fraction of neurons for hundreds to thousands of trials without any indication of neuronal damage. Metal implants may thus be exploited for stimulation of a circumscribed brain area in experimental animals, as long as RF-induced thermal changes can be kept within safe limits. Potentially, deep brain stimulation in humans could be achieved by implanting metal wires into the target areas and activating the surrounding tissue without the need of interconnects. Further research would be called for to investigate the efficacy and repeatability of such wireless stimulation approaches.

## Data and code availability

The dataset will be available from our data bank^65^ via our website: https://buzsakilab.com/wp/database/. MATLAB script packages used in the analysis of this study can be downloaded from https://github.com/MouseLand/Kilosort, and https://github.com/buzsakilab/buzcode.

## Acknowledgements

Authors thank all the members of the Buzsaki Laboratory for their support and comments on aspects of this work. This work was supported by the National Institutes of Health grant 1R01NS113782-01A1. O.Y. is supported by the NYU TL1 postdoctoral fellowship 2TL1TR001447-06A1.

## Author contributions

M.V. and O.Y. contributed equally to this work. M.V., O.Y. and G.B. conceived the project with support from L.A. and D.K.S. M.V. and O.Y. performed the experiments and associated data analysis. M.V. and G.B. wrote the paper and all authors participated in its revision. Authors declare no competing interest.

## References

1. Amon, A. & Alesch, F. Systems for deep brain stimulation: review of technical features. J. Neural Transm. 124, 1083–1091 (2017).

2. Bazaka, K. & Jacob, M. V. Implantable devices: Issues and challenges. Electron. 2, 1–34 (2012).

3. Dempsey, M. F., Condon, B. & Hadley, D. M. Investigation of the factors responsible for burns during MRI. J. Magn. Reson. Imaging 13, 627–631 (2001).

4. Dixit, N., Pauly, J. M. & Scott, G. C. Thermo-acoustic ultrasound for noninvasive temperature monitoring at lead tips during MRI. Magn. Reson. Med. 1–13 (2019) doi:10.1002/mrm.28152.

5. Erhardt, J. B. et al. Should patients with brain implants undergo MRI? J. Neural Eng. 15, (2018).

6. Finelli, D. A. et al. MR imaging-related heating of deep brain stimulation electrodes: In vitro study. Am. J. Neuroradiol. 23, 1795–1802 (2002).

7. Gupte, A. A., Shrivastava, D., Spaniol, M. A. & Abosch, A. MRI-related heating near deep brain stimulation electrodes: More data are needed. Stereotact. Funct. Neurosurg. 89, 131–140 (2011).

8. Graf, H., Steidle, G. & Schick, F. Heating of metallic implants and instruments induced by gradient switching in a 1.5-Tesla whole-body unit. J. Magn. Reson. Imaging 26, 1328–1333 (2007).

9. Dagro, A. M., Wilkerson, J. W., Thomas, T. P., Kalinosky, B. T. & Payne, J. A. Computational modeling investigation of pulsed high peak power microwaves and the potential for traumatic brain injury. Sci. Adv. 7, 1–11 (2021).

10. Wu, T., Rappaport, T. S. & Collins, C. M. The human body and millimeter-wave wireless communication systems: Interactions and implications. IEEE Int. Conf. Commun. 2015- September, 2423–2429 (2015).

11. Stijnman, P. R.. et al. Accelerating implant RF safety assessment using a low-rank inverse update. Magn. Reson. Imaging 83, 1796–1809 (2019).

12. Chou, C. K. Use of a Full-Size Human Model for Evaluating Metal Implant Heating During Magnetic Resonance Imaging. In Radio Frequency Radiation Dosimetry and Its Relationship to the Biological Effects of Electromagnetic Fields (eds. Klauenberg, B. J. & Miklavčič, D.) 473–482 (Springer Netherlands, 2000). doi:10.1007/978-94-011-4191-8_51.

13. Liam, D. The ethics of brain–computer interfaces. Nature 571, S19–S21 (2019).

14. Luechinger, R. et al. In vivo heating of pacemaker leads during magnetic resonance imaging. Eur. Heart J. 26, 376–383 (2005).

15. Ladd, M. E. & Quick, H. H. Reduction of resonant RF heating in intravascular catheters using coaxial chokes. Magn. Reson. Med. 43, 615–619 (2000).

16. Henderson, J. M. et al. Permanent neurological deficit related to magnetic resonance imaging in a patient with implanted deep brain stimulation electrodes for Parkinson’s disease: Case report. Neurosurgery 57, 1063 (2005).

17. ASTM F2503-20. Standard Practice for Marking Medical Devices and Other Items for Safety in the Magnetic Resonance Environment. 1–7 (2020).

18. Spiegel, J. et al. Transient dystonia following magnetic resonance imaging in a patient with deep brain stimulation electrodes for the treatment of Parkinson disease: Case report. J. Neurosurg. 99, 772–774 (2003).

19. Teissl, C., Kremser, C., Hochmair, E. S. & Hochmair-Desoyer, I. J. in Vitro. Radiology 3, 700–708 (1998).

20. Pavlicek, W. et al. The effects of nuclear magnetic resonance on patients with cardiac pacemakers. Radiology 147, 149–153 (1983).

21. Schenck, J. F. Safety of strong, static magnetic fields. J. Magn. Reson. Imaging 12, 2–19 (2000).

22. Winter, L., Seifert, F., Zilberti, L., Murbach, M. & Ittermann, B. MRI-Related Heating of Implants and Devices: A Review. J. Magn. Reson. Imaging 53, 1646–1665 (2021).

23. Carmichael, D. W. et al. Functional MRI with active, fully implanted, deep brain stimulation systems: Safety and experimental confounds. Neuroimage 37, 508–517 (2007).

24. Fredén Jansson, K. J., Håkansson, B., Reinfeldt, S., Taghavi, H. & Eeg-Olofsson, M. MRI induced torque and demagnetization in retention magnets for a bone conduction implant. IEEE Trans. Biomed. Eng. 61, 1887–1893 (2014).

25. Nitz, W. R. et al. On the heating of linear conductive structures as guide wires and catheters in interventional MRI. J. Magn. Reson. Imaging 13, 105–114 (2001).

26. Nordbeck, P. et al. Measuring RF-induced currents inside implants: Impact of device configuration on MRI safety of cardiac pacemaker leads. Magnetic Resonance in Medicine vol. 61 570–578 (2009).

27. Calcagnini, G. et al. In vitro investigation of pacemaker lead heating induced by magnetic resonance imaging: Role of implant geometry. J. Magn. Reson. Imaging 28, 879–886 (2008).

28. Babouri, A. & Hedjeidj, A. In vitro investigation of eddy current effect on pacemaker operation generated by low frequency magnetic field. In Annual International Conference of the IEEE Engineering in Medicine and Biology Society. IEEE Engineering in Medicine and Biology Society. 5684–5687 (2007).

29. Krupa, K. & Bekiesińska-Figatowska, M. Artifacts in magnetic resonance imaging. Polish J. Radiol. 80, 93–106 (2015).

30. Boutet, A. et al. Improving Safety of MRI in Patients with Deep Brain Stimulation Devices. Radiology 296, 250–262 (2020).

31. ISO/TS 10974:2018 Assessment of the safety of magnetic resonance imaging for patients with an active implantable medical device. (2018).

32. Voroslakos, M. et al. 3D-printed Recoverable Microdrive and Base Plate System for Rodent Electrophysiology. Bio-Protocol vol. 11 (2021).

33. Vöröslakos, M., Petersen, P. C., Vöröslakos, B. & Buzsáki, G. Metal microdrive and head cap system for silicon probe recovery in freely moving rodent. Elife 10, 1–21 (2021).

34. Moser, E., Mathiesen, I. & Andersen, P. Association between brain temperature and dentate field potentials in exploring and swimming rats. Science (80-.). 259, 1324–1326 (1993).

35. Petersen, P. C., Voroslakos, M. & Buzsáki, G. Brain temperature affects quantitative features of hippocampal sharp wave ripples. J. Neurophysiol. 127, 1417–1425 (2022).

36. Yaghmazadeh, O. et al. Neuronal activity under transcranial radio-frequency stimulation in metal-free rodent brains in-vivo. Commun. Eng. 1–15 (2022) doi:10.1038/s44172-022-00014-7.

37. Pachitariu, M., Steinmetz, N., Kadir, S., Carandini, M. & Harris, K. D. Kilosort: realtime spike-sorting for extracellular electrophysiology with hundreds of channels. bioRxiv 061481 (2016).

38. Schomburg, E. W. et al. Theta Phase Segregation of Input-Specific Gamma Patterns in Entorhinal-Hippocampal Networks. Neuron 84, 470–485 (2014).

39. Watson, B. O., Levenstein, D., Greene, J. P., Gelinas, J. N. & Buzsáki, G. Network Homeostasis and State Dynamics of Neocortical Sleep. Neuron 90, 839–852 (2016).

40. Racine, R. J. Modification of seizure activity by electrical stimulation: II. Motor seizure. Electroencephalogr. Clin. Neurophysiol. 32, 281–294 (1972).

41. Lynch, M. & Sutula, T. Recurrent excitatory connectivity in the dentate gyrus of kindled and kainic acid-treated rats. J. Neurophysiol. 83, 693–704 (2000).

42. Foster, K. R. & Morrissey, J. J. Thermal aspects of exposure to radiofrequency energy: Report of a workshop. Int. J. Hyperth. 27, 307–319 (2011).

43. Rahimpour, S., Kiyani, M., Hodges, S. E. & Turner, D. A. Deep brain stimulation and electromagnetic interference Shervin. Clin. Neurol. Neurosurg. 203, 139–148 (2021).

44. Sidiropoulos, C. et al. Intraoperative MRI for deep brain stimulation lead placement in Parkinson’s disease: 1 year motor and neuropsychological outcomes. J. Neurol. 263, 1226–1231 (2016).

45. Golestanirad, L. et al. Reducing RF-induced Heating near Implanted Leads through High-Dielectric Capacitive Bleeding of Current (CBLOC). IEEE Trans. Microw. Theory Tech. 67, 1265–1273 (2019).

46. Vöröslakos, M. et al. Direct effects of transcranial electric stimulation on brain circuits in rats and humans. Nat. Commun. 9, 483 (2018).

47. Ozen, S. et al. Transcranial electric stimulation entrains cortical neuronal populations in rats. J. Neurosci. 30, 11476–85 (2010).

48. El Khoueiry, C. et al. Decreased spontaneous electrical activity in neuronal networks exposed to radiofrequency 1,800 mhz signals. J. Neurophysiol. 120, 2719–2729 (2018).

49. Moretti, D. et al. In-vitro exposure of neuronal networks to the GSM-1800 signal. Bioelectromagnetics 34, 571–578 (2013).

50. Tattersall, J. E.. et al. Effects of low intensity radiofrequency electromagnetic fields on electrical activity in rat hippocampal slices. Brain Res. 904, 43–53 (2001).

51. Thuroczy, G., Kubinyi, G., Bakos, J., Szabo, L. D. & Bodo, M. Simultaneous response of brain electrical activity (EEG) and cerebral circulation (REG) to microwave exposure in rats. Rev. Environ. Health 10, 135–148 (1994).

52. Beason, R. C. & Semm, P. Responses of neurons to an amplitude modulated microwave stimulus. Neurosci. Lett. 333, 175–178 (2002).

53. Eulitz, C., Ullsperger, P., Freude, G. & Elbert, T. Mobile phones modulate response patterns of human brain activity. Neuroreport 9, 3229–3232 (1998).

54. Ghosn, R. et al. Radiofrequency signal affects alpha band in resting electroencephalogram. J. Neurophysiol. 113, 2753–2759 (2015).

55. Vecsei, Z. et al. Short-term radiofrequency exposure from new generation mobile phones reduces EEG alpha power with no effects on cognitive performance. Sci. Rep. 8, 1–12 (2018).

56. Kiyatkin, E. A., Brown, P. L. & Wise, R. A. Brain temperature fluctuation: A reflection of functional neural activation. Eur. J. Neurosci. 16, 164–168 (2002).

57. Guidelines for Evaluating the Environmental Effects of Radiofrequency Radiation. FCC- 96–326 (1996).

58. Chou, C. K., Guy, A. W. & Galambos, R. Auditory perception of radio-frequency electromagnetic fields. J. Acoust. Soc. Am. 71, 1321–1334 (1982).

59. Chou, C.-K & Guy, A. W. Microwave-induced auditory responses in guinea pigs: Relationship of threshold and microwave-pulse duration. Radio Sci. 14, 193–197 (1979).

60. Lin, J. C. & Wang, Z. Hearing Of Microwave Pulses By Humans And Animals: Effects, Mechanism, And Thresholds. Health Phys. 92, 621–628 (2007).

61. Tagliati, M. et al. Safety of MRI in patients with implanted deep brain stimulation devices. Neuroimage 47, T53–T57 (2009).

62. Shrivastava, D. et al. Heating Induced near Deep Brain Stimulation Lead Electrodes during Magnetic Resonance Imaging with a 3T Transceive Volume Head Coil. Phys. Med. Biol. 57, 5651–5665 (2012).

63. Boutet, A. et al. 3-Tesla MRI of deep brain stimulation patients: safety assessment of coils and pulse sequences. J. Neurosurg. 132, 586–594 (2020).

64. Musk, E. & Neuralink. An integrated brain-machine interface platform with thousands of channels. J. Med. Internet Res. 21, 1–14 (2019).

65. Petersen, P. C., Hernandez, M. & Buzsáki, G. The Buzsaki Lab Databank - Public electrophysiological datasets from awake animals. Zenodo https://zenodo.org/record/4307883 (2020) doi: https://doi.org/10.5281/zenodo.4307883.

