## Supplementary material for "Brain-implanted conductors amplify radiofrequency fields in rodents: advantages and risks"

György Buzsáki

Department of Neurology, Grossman School of Medicine, New York University, New York, NY 10016, USA

Leeor Alon and Daniel K. Sodickson

Department of Radiology, Grossman School of Medicine, New York University, New York, NY 10016, USA

György Buzsáki

| **Source field** | **Interaction** | **Hazard** |
| --- | --- | --- |
| RF | Induced RF Heating | Focused RF tissue heating near device |
|  | Electromagnetic induction | Induced voltages at implantable medical device’s terminals, rectification, induced stimulation |

**Table 1. Source of electromagnetic field and related hazards for active implantable medical devices**^17^**.**

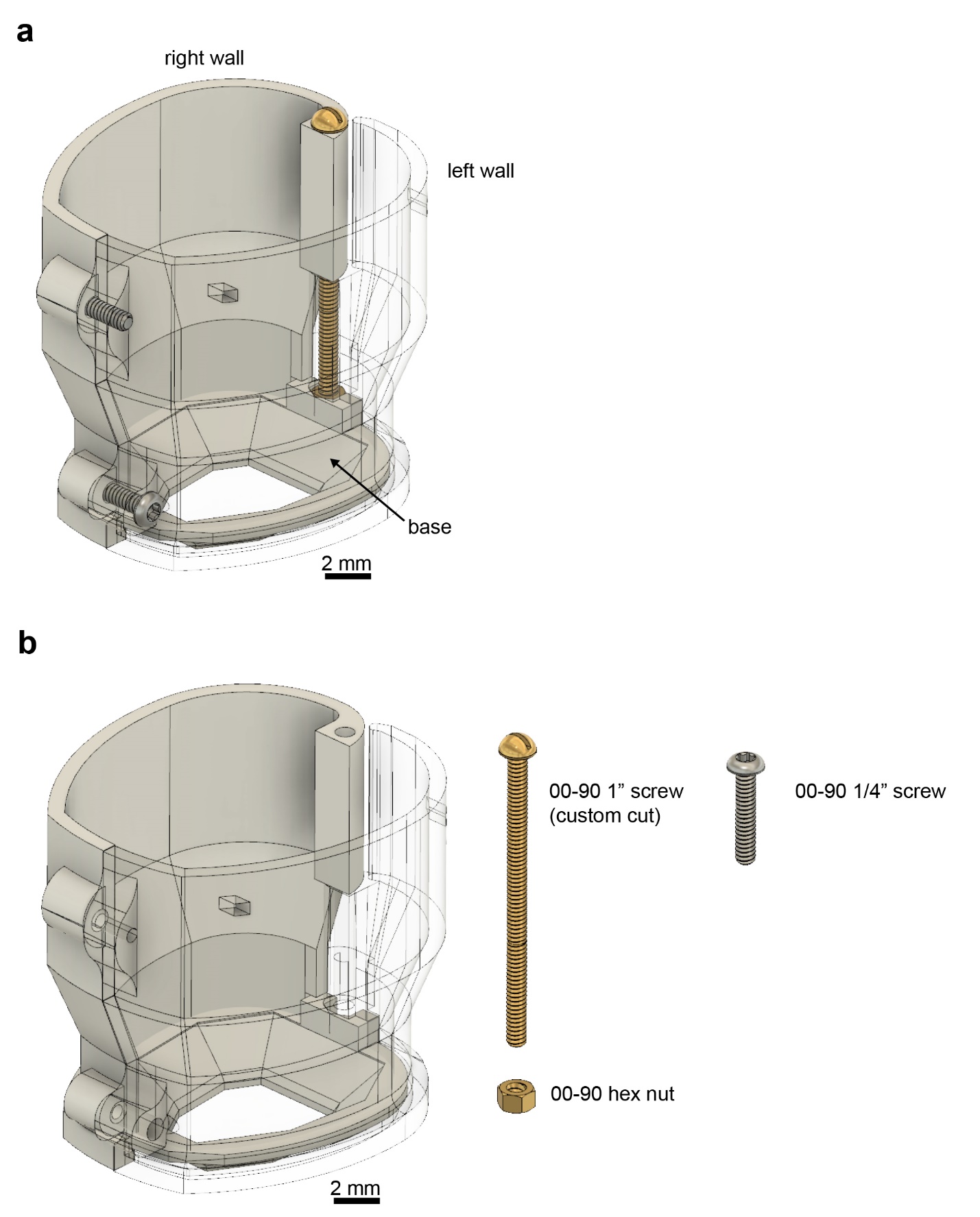

**Suppl. Figure 1. Modular plastic head cap enables step-by-step surgery procedure.** **a** Fully assembled head cap (the weight is 1.895g). The head cap consists of a base, a left and a right wall. Note that left wall is displayed as transparent to improve visibility of other components. **b** Components of the head cap. A hex nut (00-90 size) is glued to the base. A custom-cut 1” screw holds the walls in the back and two ¼” screws secure the front of the cap (all screws are 00-90 size). For further details, see Supplementary video 1.

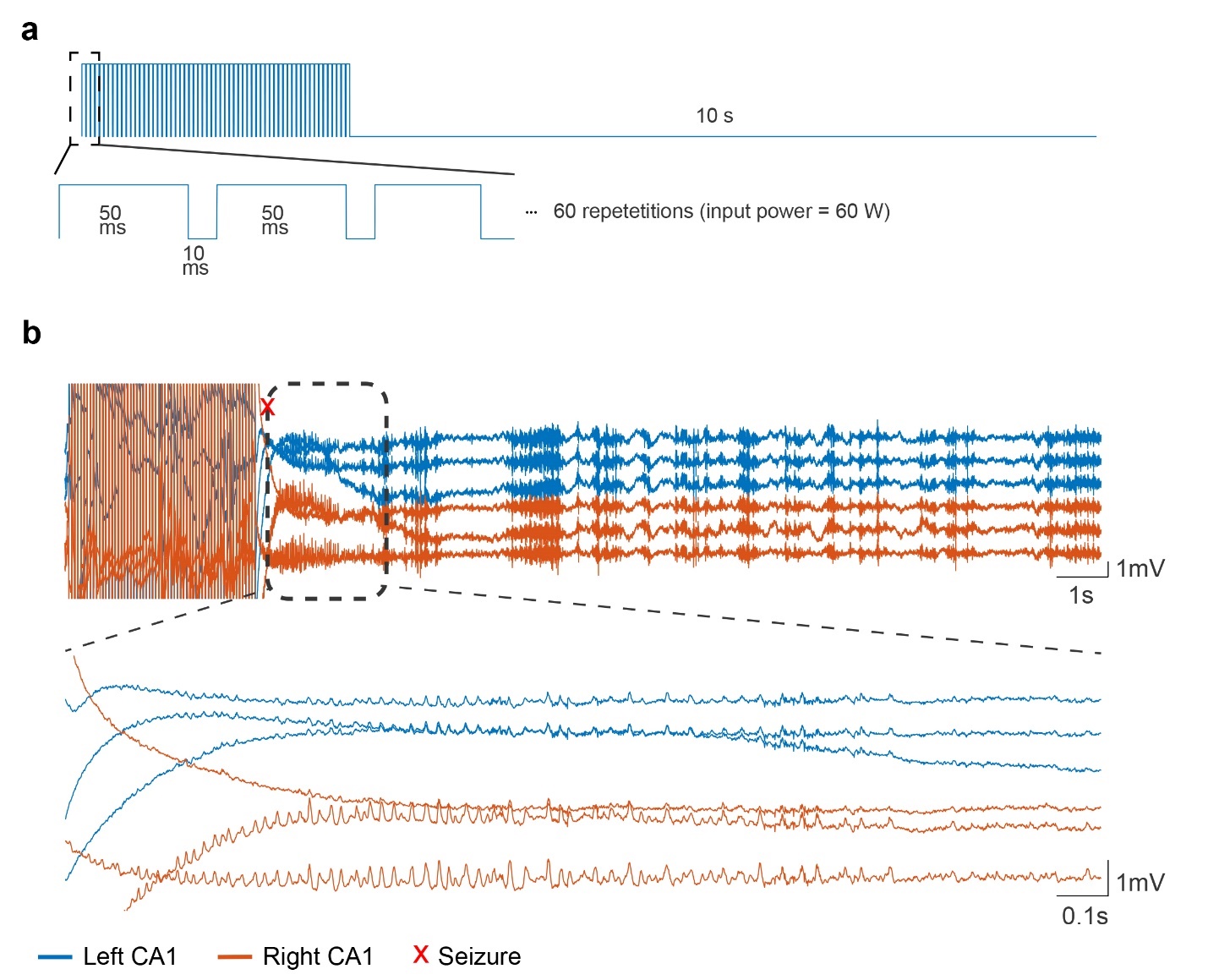

**Suppl. Figure 2. Repetition frequency of RF stimulation did not affect the seizure occurrence**. **a** RF-stimulation was applied using a patch antenna (50 ms pulses at 100 Hz, 60 repetitions, 10 s off period between stimulations, input power was 60 W). **b** RF stimulation-induced seizure in a mouse implanted with bilateral tungsten triplet recording electrodes. Bottom panel shows a zoomed-in portion of the signal. Note that stimulation induced bilateral afterdischarge in the hippocampi.

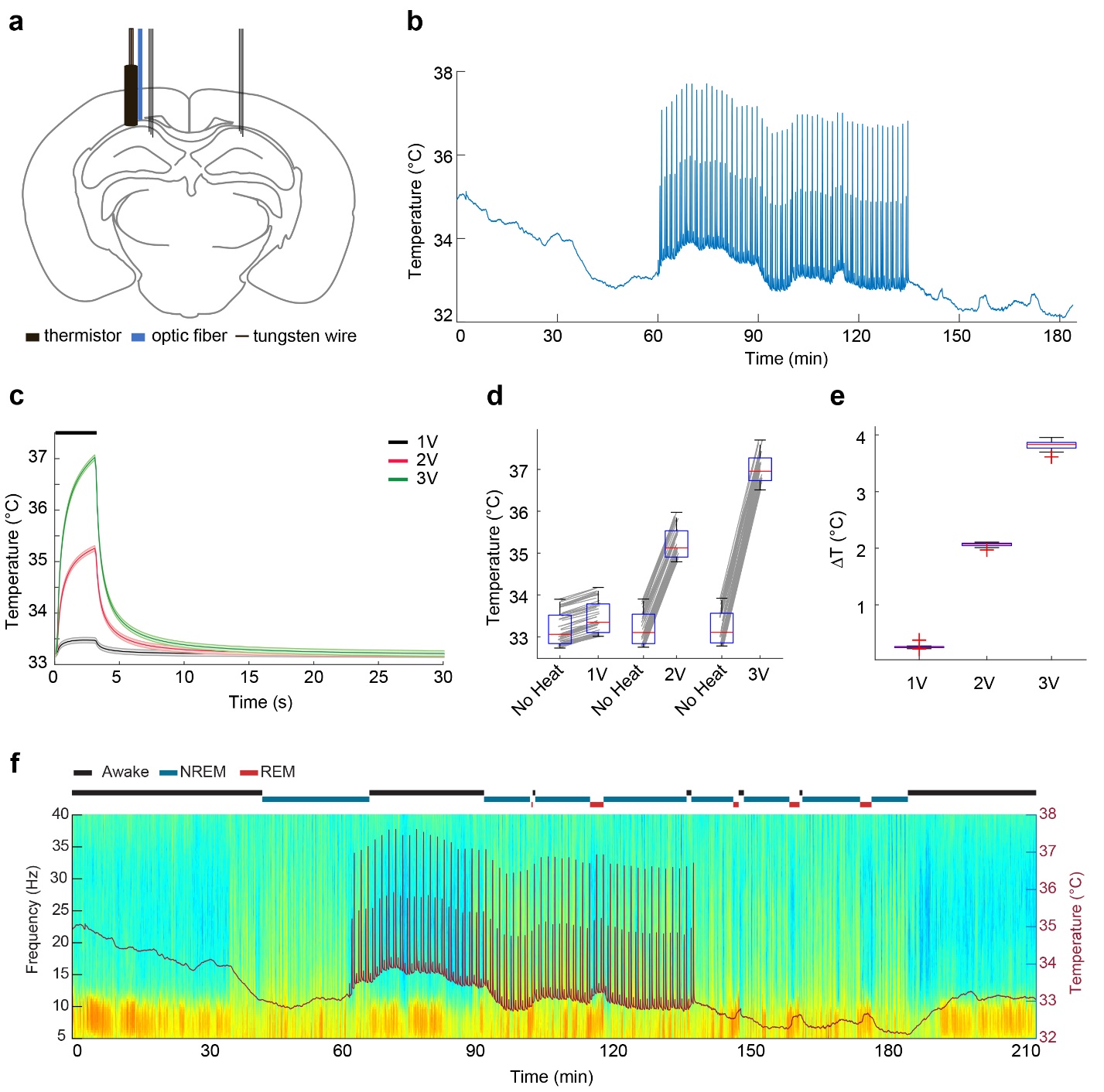

**Suppl. Figure 3. Local heating of the hippocampus within the physiological range does not generate afterdischarges**. **a** Mice were implanted with an optic fiber attached to a thermistor and 2 tungsten triplet recording electrodes. **b** Physiological brain temperature variation over time and induced temperature rise due to optical heating. **c-e** Three seconds of optical heating can increase brain temperature reliably (black bar represents the time of heating). Three different optical powers were applied (n = 50 trials, 3 seconds heating, 27 seconds no heating). **C** shows mean±SEM, **d** shows the trial-by-trial variations and **e** shows the absolute change in temperature during heating. **f** Time-power analysis of hippocampal local field potentials (LFPs) and brain temperature (orange line overlaid on spectrogram). Single channel LFP from the hippocampal CA1 pyramidal layer was used to calculate the time-resolved fast Fourier transform-based power spectrum. Automatic brain state scoring is shown above the spectrum (Wake, NREM and REM, black, blue, and red lines).

**Suppl. Video 1. RF stimulation induced effects in mice implanted with recording electrode**

**Suppl. Video 2. RF stimulation induced effects in mice without brain implants**
